# Effects of high pCO_2_ on snow crab embryos: Ocean acidification does not affect embryo development or larval hatching

**DOI:** 10.1101/2022.10.06.511099

**Authors:** W. Christopher Long, Katherine M. Swiney, Robert J. Foy

**Affiliations:** Kodiak Laboratory, Alaska Fisheries Science Center, National Marine Fisheries Service, National Oceanic and Atmospheric Administration

## Abstract

Ocean acidification, a decrease in ocean pH due to absorption of anthropogenic CO_2_, has variable effects on different species. To examine the effects of decreased pH on snow crab (*Chionoecetes opilio*), a commercial species in Alaska, we reared ovigerous females in one of three treatments: ambient pH (~8.1), pH 7.8, and pH 7.5, through two annual reproductive cycles. Morphometric changes during development and hatching success were measured for embryos both years and calcification was measured for the adult females at the end of the 2-year experiment. Embryos and larvae analyzed in year one were from oocytes developed, fertilized, and extruded *in situ*, whereas embryos and larvae in year two were from oocytes developed, fertilized, and extruded under acidified conditions in the laboratory. Embryo morphology during development was unaffected by pH during both years. The number of successfully hatched live larvae was unaffected by pH treatment in both years. Embryo mortality was very low, hatching success high, and neither differed with treatment in either year. Percent calcium in adult females’ carapaces did not differ among treatments at the end of the experiment. The results from this two-year study suggest that snow crabs are well adapted to projected ocean pH levels within the next 2 centuries, although other life-history stages still need to be examined for sensitivity.

## Introduction

The anthropogenic release of CO_2_ is causing an increase in atmospheric pCO_2_ which then dissolves into the oceans. The increased pCO_2_ in the oceans changes the carbonate chemistry, decreasing the pH and the saturation state of calcium content [1] in a process known as ocean acidification (OA). Since the beginning of the industrial revolution the pH of surface oceanic waters has dropped by approximately 0.1 units with a further ~0.3 unit reduction predicted by the end of this century [1–3]. High latitude waters are likely to acidify more rapidly than elsewhere, in part because CO_2_ is more soluble in colder waters and in part because of the upwelling of deep water rich in CO_2_ [4]. This change in ocean chemistry may have deleterious effects on many marine organisms, and, because of the accompanying decrease in calcium carbonate saturation state, calcifying marine organisms may be particularly vulnerable [5]. There is a great deal of variability both within and among marine taxa in their response to OA [6,7] and this variability is projected to alter marine food webs and ecosystems [8].

Decapod crustaceans, many of which are valuable commercial species, are a calcifying marine taxa that can be strongly effected by OA. When exposed to low pH many decapods respond through active ion transport in the gills, typically either H^+^/Na^+^ or Cl^-^/HCO3^-^ exchange, to maintain acid/base homoeostasis in their hemolymph [9], although this is not effective in all species or life history stages [10]. However, this active transport comes at an energetic cost which may divert energy away from other processes; this and other effects of OA may result in a range of effects including reduced hatching success [11,12], slower growth [13–15], higher mortality [16–18], deformities [19,20], immune disruption or increased disease rate [21,22], reduced calcium content or hardness in the exoskeleton [17,23,24], and altered behavior [25,26] in many species. However, other species are highly resilient to OA [27,28]. For species that are negatively affected, reductions in ocean pH are predicted to result in decreases in population size, and for species that are commercially harvested, the fisheries that depend on them [29–31].

Not only does OA have direct effects on marine species, but carryover effects, where exposure at one life history stage affects fitness in a subsequent life history stage, frequently occur, although they are not as frequently studied. For example, for both red king crab, *Paralithodes camtschaticus*, and the spider crab *Hyas araneus* embryo exposure to high pCO_2_ reduces survival at the larval stage [32,33]. Maternal effect, a sub-type of transgenerational effects, are when exposure of the mother during oocyte development affects larval or juvenile performance and are a particular sub-set of carryover effects [34]. Sometimes, as in the case of the muscle *Musculista senhousia*, maternal effects can be positive [35] indicating adaptive potential, but in other species maternal effects can be negative [12] suggesting reduced maternal investment in oocytes under stressful conditions. Therefore, it is necessary to quantify if and to what degree carryover effects occur for a particular species to avoid over or underestimating the effects of OA.

Snow crabs, *Chionoecetes opilio*, are a high latitude species with populations ranging across the North Pacific from Japan to the United States and up into the Arctic in the Beaufort and Chukchi Seas, another in the Gulf of St. Lawrence, and another in the Barents Sea, where it is an invasive species [36,37]. It is also an important fisheries species; in 2018 the ex-vessel value was $75.2 million in the Bering Sea fishery alone [38]. Across its range, even where it is not fished, snow crab can be a dominant biomass in the benthic community and play an important role as a primary/secondary benthic predator within the ecosystem [39]. Snow crab have a life history that is typical of many decapod crustaceans. Pubescent females undergo a terminal molt to maturity and mate before extruding a clutch of eggs in the early spring [40]. Eggs are brooded for either approximately a year or two years (depending on temperature) and hatch in the late spring [41–44]. The larvae are planktivorous and pass through two zoeal and one megalops stage before settling to the benthos and molting to the first crab stage [45]. Although no work has been done to date examining the effects of OA on snow crab, a congener species, the southern Tanner crab, *C. bairdi*, is highly vulnerable to OA; OA can reduce hatching success by over 70% [12], affects larval fitness, and decreases juvenile growth and survival [46], although the larvae are fairly resilient [47]. Given its high value as a commercial species, its importance to benthic ecosystems, and the vulnerability of closely related species, it is an important target species for OA research. In this paper, we present the results of a study to determine both the direct and maternal effects of OA on snow crab embryos and fecundity.

## Methods

### Overview

Female crabs in this experiment were held through two brooding/hatching cycles. In the first year, wild-extruded embryos were exposed to three pH treatments to quantify the direct effects of OA on embryo development. At hatching, larvae were collected individually from each female and hatching success and fecundity were determined. After hatching, females were allowed to mate with a male, extruded a new clutch of eggs, and were held in the same treatment pH as the first year throughout embryo development and larval hatching. The second year allowed us to examine the combination of maternal (or carryover) effects and direct effects on embryo development and hatching success.

### Female collection and holding

Female snow crab with newly-extruded uneyed eggs and mature male snow crabs were captured on the 2014 eastern Bering Sea trawl survey [48] and transported to the Kodiak Fisheries Research Center in coolers packed with damp burlap and ice packs via airplane from Dutch Harbor on July 16, 2014. Females were held in tanks with flow-through sea water at ambient pH and salinity chilled to 2°C briefly before the beginning of the experiment on August 6, 2014; males were held in the same conditions (ambient pH) until needed for mating (see below). Sand filtered seawater at ambient salinity from Trident Basin intakes at 15 and 26m was used throughout this experiment. Throughout holding and the entire experimental period crabs were fed a diet of frozen herring and squid to excess twice a week. Twenty five crabs were randomly assigned to each of one of three pH treatments based on projected future ocean pH levels: current surface ambient (~pH 8.1), pH 7.8 (projected for 2100), and pH 7.5 (projected for 2200). Only healthy crabs with full clutches and no more than 2 missing limbs were used. During year 1, 15 randomly selected crabs from each treatment were sampled for embryo development and hatching (see below). During year two, 14 females, comprised of the crabs from the original 15 that had survived the first year, plus additional crabs from the additional crab held were used in each treatment. Females that were not selected for sampling were held in the same manner as those which were.

During the majority of the experiment, crabs within each treatment group were held communally in experimental holding tanks, one per treatment, (0.6 m x 1.2m x 0.6m) supplied with flow-through water at the appropriate pH (see below for details on water acidification). A temperature of 2°C was maintained in each tank with a recirculating chiller. Both years, as crabs neared the time for hatching they were transferred to individual 68 L tubs. This was done so that hatching data could be collected for each individual female. All hatching tubs received recirculating water from a 2000 L head tank (one per treatment) that received flow-through seawater, was adjusted to the appropriate pH (see below), and maintained at 2°C with a recirculating chiller. Although this design does not allow us to determine if there was a tank effect as crab were held either together or received water from a communal head tank, the fact that the embryos are held under the abdominal flap and protected by each female is sufficient isolation to treat each as an independent replicate. In the first year, after a female stripped her pleopods (cleaned off the empty egg cases and remaining unhatched eggs in preparation for extruding a new clutch), she was held with a male as a potential mate to ensure there was no potential for sperm limitation in the second year; however, as female snow crab can store sperm we were unable to determine if a female was mated and whether fresh sperm from that mating, stored sperm, or a combination of stored and fresh sperm was used to fertilize her new clutch. Males were randomly assigned to females and no male was used as a potential mate for more than two females. After all the females had extruded a new clutch of eggs, they were transferred back to communal holding in the experimental holding tanks.

### Water acidification and chemistry

In both the experimental tanks and the hatching tubs the seawater was acidified using CO_2_. The water flowing into the holding tanks was acidified as described in [46]. In brief, water was acidified down to ~pH 5.5 by bubbling CO_2_. This water was mixed with seawater in head tanks using peristaltic pumps controlled with Honeywell controllers and Duafet III pH probes in the head tanks to achieve the nominal pH. Water from the head tanks then flowed into the experimental holding tanks. When crabs were transferred to the hatching tubs, the head tank supplying the water was acidified by bubbling CO_2_, the flow of which was controlled by Honeywell controllers and Durafet III pH probe in the head tanks. Temperature and pH (free scale) were measured daily in experimental units using a Durafet III pH probe calibrated with TRIS buffer [49]. Water from the head tanks was sampled once per week (N = 98 per treatment) and samples were poisoned with mercuric chloride and analyzed for dissolved inorganic carbon (DIC) and total alkalinity (TA) at an analytical laboratory. DIC and TA were determined using a VINDTA 3C (Marianda, Kiel, Germany) coupled with a 5012 Coulometer (UIC Inc.) according to the procedure in [50] using Certified Reference Material from the Dickson Laboratory (Scripps Institute, San Diego, CA, USA; [51]). The other components of the carbonate system were calculated in R (V3.6.1, Vienna, Austria) using the seacarb package [52].

Treatment pHs for the two acidified treatments exactly averaged the target levels across two years of treatment and water temperatures were well maintained throughout the experiment (Table 1). The ambient pH treatment was supersaturated with regards to both calcite and aragonite while the pH 7.8 treatment was undersaturated with regards to aragonite and the pH 7.5 treatment was undersaturated with regards to both (Table 1).

**Table 1.** Water physical and chemical parameters.

| Treatment | Temperature | pH<sub>F</sub> | pCO<sub>2</sub><br>μatm | HCO<sub>3</sub><sup>-</sup><br>mmol/kg | CO<sub>3</sub><sup>-2</sup><br>mmol/kg | DIC<br>mmol/kg | Alkalinity<br>mmol/kg | Ω<sub>Aragonite</sub> | Ω<sub>Calcite</sub> |
| --- | --- | --- | --- | --- | --- | --- | --- | --- | --- |
| Ambient | 2.09 ± 0.32 | 8.11 ± 0.08 | 362.18 ± 68.33 | 1.90 ± 0.05 | 0.09 ± 0.02 | 2.01 ± 0.04 | 2.11 ± 0.02 | 1.36 ± 0.23 | 2.19 ± 0.37 |
| pH 7.8 | 1.97 ± 0.30 | 7.80 ± 0.02 | 760.98 ± 43.95 | 2.00 ± 0.04 | 0.05 ± 0.00 | 2.09 ± 0.05 | 2.09 ± 0.21 | 0.69 ± 0.04 | 1.11 ± 0.06 |
| pH 7.5 | 2.05 ± 0.31 | 7.50 ± 0.02 | 1548.29 ± 102.11 | 2.04 ± 0.06 | 0.02 ± 0.00 | 2.15 ± 0.06 | 2.11 ± 0.02 | 0.36 ± 0.02 | 0.57 ± 0.04 |

Water parameters in experimental tanks during snow crab experiments. Temperature and pH were measured daily, dissolved
inorganic carbon (DIC) and alkalinity were measured weekly, and all other parameters were calculated. Values are mean ± 1 SD.

### Embryo development

Embryo development was monitored throughout both years. Once a month, approximately 20 eggs were randomly sampled from each female. The embryonic developmental stage was determined per Moriyasu and Lanteigne [42]. Uneyed eggs were stained for 5 minutes with Bonin’s solution to facilitate observation of the external morphology of the embryos; eyed eggs were not stained. The embryonic stages were determined under a stereo microscope at ~63x magnification. Additionally, digital images of ten fresh eggs from each female were taken with a digital camera attached to the same microscope at a total magnification of 63x. Measurements were made as per Swiney et al. [12]. Using image analysis software (Image Pro Plus Version 7.0.1.658, Media Cybernetics, Inc., Rockville, Maryland USA), egg area and diameters (maximum, minimum, and average) were measured. Once embryos were discernable, embryo areas and yolk areas and diameters (maximum, minimum, and average) were also measured. Lastly, when embryos become eyed, eyespot area and diameters (maximum, minimum, and average) were measured. Percent area yolk (PAY) was calculated as: 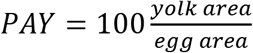. For each female, within each sampling period, the 10 measured embryos were averaged prior to analysis. In May 2015, all but five of the females had already extruded new clutches of eggs; as there were so few females, measurements made in May on their embryos were not included in either the analysis of embryo stage or embryo morphometrics.

Embryo stage was analyzed separately for each year using an ANOVA with pH treatment fully crossed with month and female (nested within treatment) as factors. Embryo morphometrics were analyzed separately for each year and normalized (expressed in terms of their z-score) prior to analysis. Morphometrics were analyzed using permutational MANOVA (PERMANOVA) on a Euclidian distance resemblance matrix with pH treatment fully crossed with month and female (nested within treatment) as factors. Data were visualized using a non-metric multidimensional scaling plot (MDS).

### Fecundity, hatching success, and mineral content

Before hatching began, crabs were moved into individual tubs and larvae were collected in a net at the outflow of each tub. Each day, every net was checked and the number of larvae estimated. If there were fewer than ~100 larvae the larvae were counted; if there were more than 100, the number hatched was estimating using dry mass. The larvae were collected, rinsed briefly in DI water, dried to a constant mass at 60°C and weighed. For each female, five times throughout hatching a subset of 50 larvae were counted out, dried to a constant mass, and the dry mass was determined. The average larval mass for each female was calculated and use to estimate the total number hatched each day. On a number of occasions, larvae were collected live for experiments (results from the larval experiments are reported separately [53]) and could not be dried. In these cases, the number of larvae was estimated by taking 3-4 subsamples of known volume, counting the number of larvae in each, and using the average concentration of larvae to calculate the total number.

Fecundity was defined as the number of viable larvae hatched. Hatching success was defined as the percent of the total estimated number of embryos (viable larvae + non-viable larvae + unhatched eggs) that hatched into viable larvae. Non-viable larvae were defined as larvae which hatched but failed to molt from the pre-zoeal to the first zoeal stage. After hatching was complete and females had stripped their pleopods, the debris was collected and examined microscopically and the total number of unhatched eggs, and viable and non-viable larvae were estimated from volumetric subsamples. The percent non-viable larvae and unhatched eggs were also calculated for each female. At the end of the final year all females were sacrificed and the calcium and magnesium content in their exoskeletons was determined in an analytical laboratory from an approximately 2 x 2 cm sample of the carapace taken from the posterior margin.

Fecundity, hatching success, percent non-viable larvae, percent unhatched eggs, and carapace calcium and magnesium contents were all analyzed with a one-way ANOVA with pH treatment as the factor.

## Results

### Data availability

All raw data from these experiments and the associated metadata are available at the National Centers for Environmental Information [54].

### Embryo development

The average embryo stage increased over time in both years of the study but did not differ among pH treatment (Fig. 1). Stage differed among months in both years (Year 1: F_8,227_ = 2.412, p < 0.0005; Year 2: F_11,326_ = 1,141.6, p > 0.0005) but not among treatments (Year 1: F_2,227_ = 2.412, p = 0.092; Year 2: F_2,326_ = 1.097, p = 0.335). The interaction between month and treatment was significant in the second, but not the first year (Year 1: F_16,227_ = 0.535, p = 0.928; Year 2: F_22,326_ = 2.838, p < 0.0005); however, in the second year, there was no significant difference among the treatments within any single month (Bonferoni’s post-hoc test, minimum p = 0.087).

**Fig. 1.**
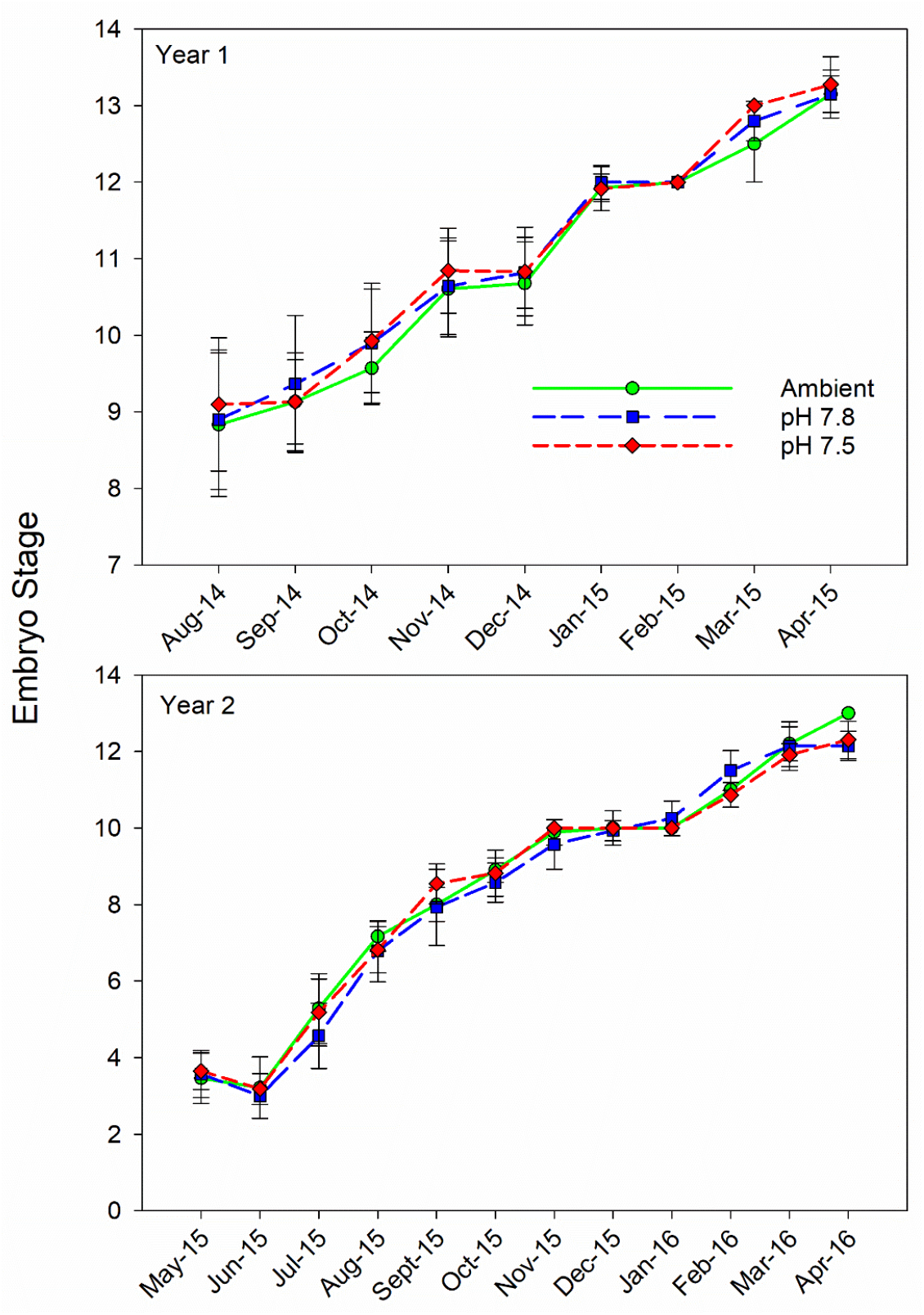
Effect of pH on snow crab embryo stage. Effects of three different pH treatments on the stage of embryo development in snow crabs over two successive brooding cycles (Year 1 and Year 2). Symbols are the mean stage for each month and error bars are one standard deviation.

Embryo morphometrics showed a clear progression in development among months (Fig. 2) in both year 1 (PERMANOVA; Pseudo-F_8,272_ = 467.63, p = 0.001) and year 2 (Pseudo-F_11,327_= 549.29, p = 0.001). There was no difference among treatments in either year 1 (Pseudo-F_2,272_ = 1.065, p = 0.385) or year 2 (Pseudo-F_2,327_= 549.29, p = 0.469), but there was a significant interactive effect in both years (year 1: Pseudo-F_16,272_ = 1.451, p = 0.001; year 2: Pseudo-F_22,327_ = 1.6146, p = 0.003); however, there was no clear pattern discernable. In the first year, a post-hoc PERMANOVA indicated a significant difference between pH 7.8 and ambient embryos in October and a difference between pH 7.8 and the other two treatments in April; in the second year there was a significant difference between ambient and pH 7.5 embryos in December but no other month. Given large number of contrasts (60 total), the lack of any pattern either within or between years, and the strong overlap of the treatments in the MDS plots in both years, we interpret these differences as representing type I statistical errors.

**Figure 2.**
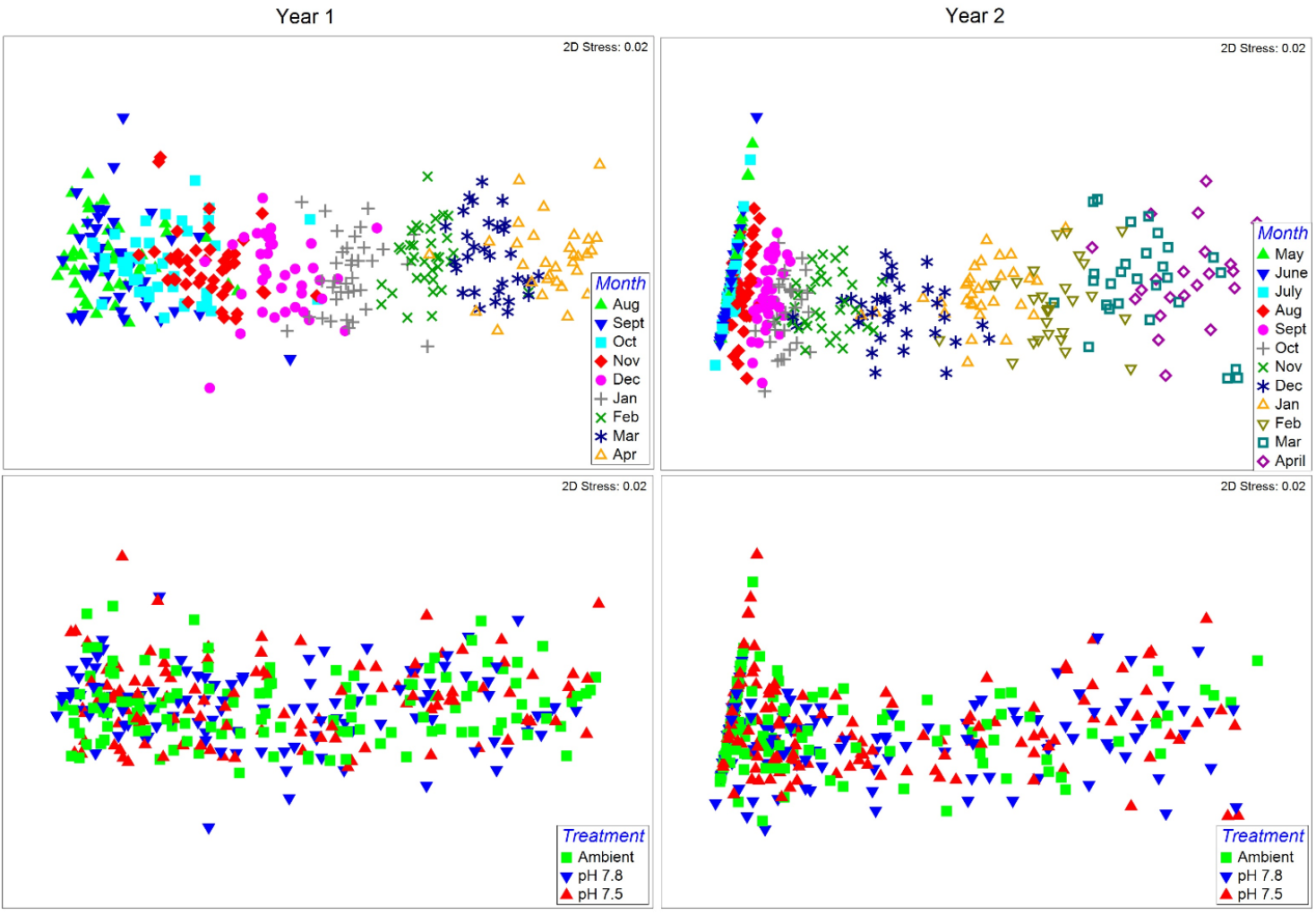
Effect of pH on snow crab embryo morphometrics. Non-metric multidimensional scaling plots of snow crab embryo morphometrics measured monthly during two brooding cycles (years) and reared in 3 different pH treatments. Plots on the left represent the year 1 embryos (first brooding cycle), and those on the right represent year 2 embryos. Top row of plots graph the data by month to show embryo development and the bottom 2 rows graph the data by pH treatment.

### Fecundity, hatching success, and mineral content

Fecundity was higher in the second year but did not differ among treatments in either the first or second year (Fig. 3, Table 2). Hatching success was high during both years and did not differ among treatments; in year 1 hatching success was greater than 95% in all treatments and in year 2 it was greater than 98% (Fig. 4, Table 2). Similarity the percent of non-viable larvae never averaged above 2% for any treatment and the percent of unhatched eggs was never above 4% (Fig 4, Table 2). The percent Ca and Mg and the Ca:Mg in the female carapaces did not differ among treatments at the end of the experiment (Fig. 5, Table 2).

**Fig. 3.**
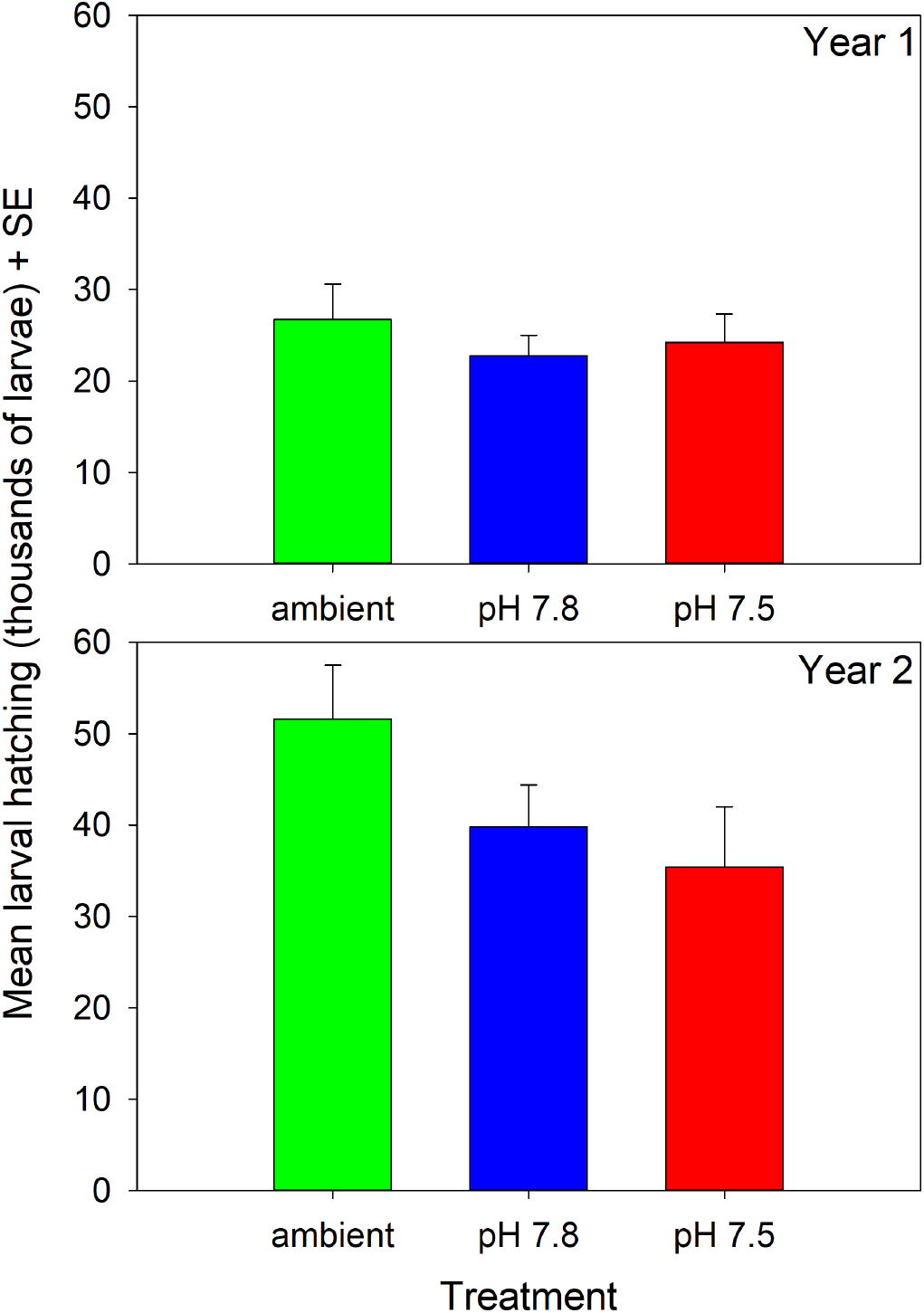
Effect of pH on snow crab fecundity. Fecundity, defined as the number of viable larvae hatched, in females held in 3 different pH treatments over 2 years. There are no statistically significant differences among treatments in either year.

**Fig. 4.**
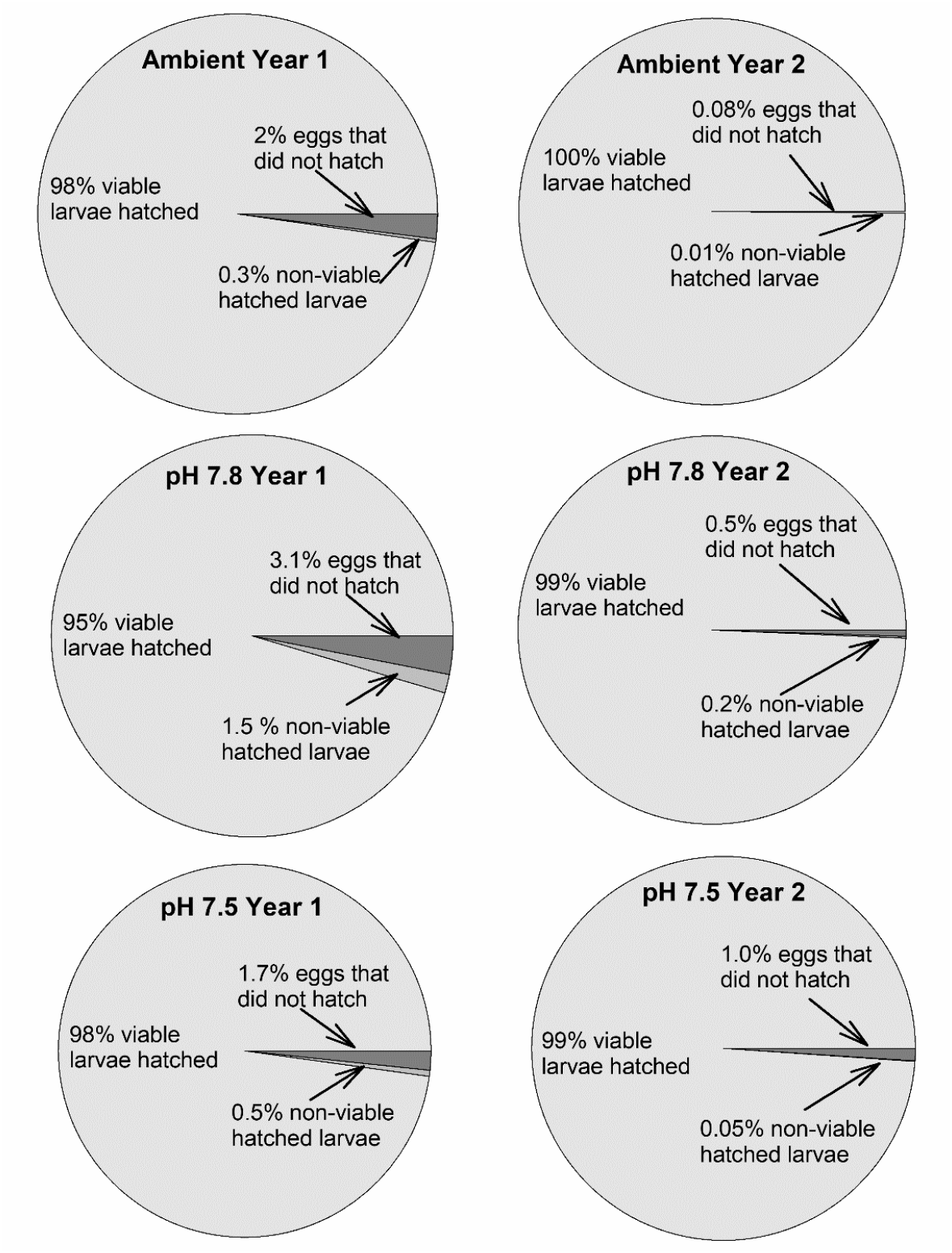
Effect of pH on snow crab hatching success. Hatching success in female snow crabs held in three different pH treatments for 2 years. There are no statistically significant differences among treatments in either year.

**Figure 5.**
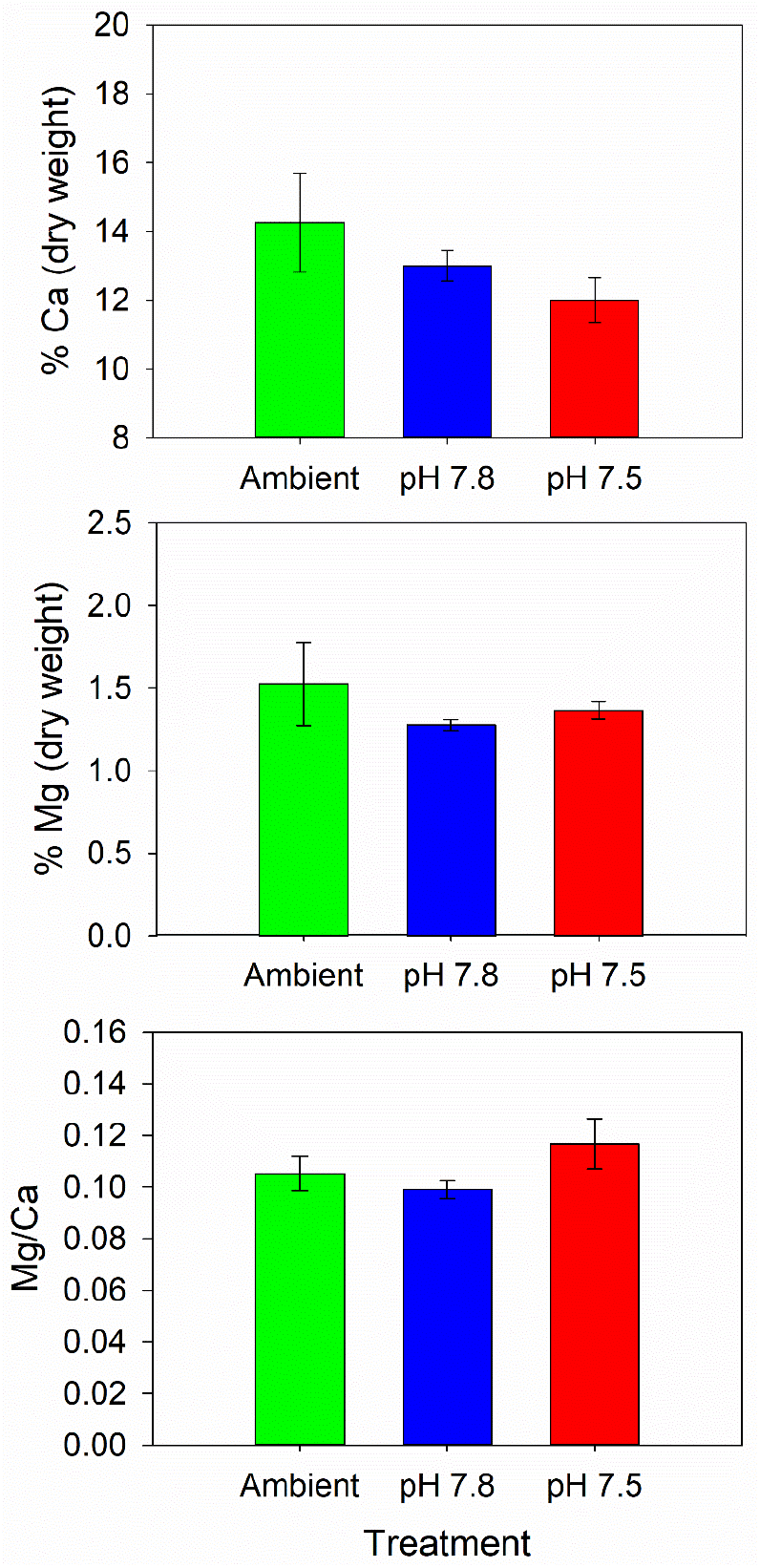
Effect of pH on snow crab carapace mineral content. Calcium and magnesium content and Mg:Ca ratio in the carapaces of female snow crabs held for 2 years in 3 different pH treatments. Error bars are 1 standard deviation. There are no statistically significant differences among treatments.

**Table 2.**
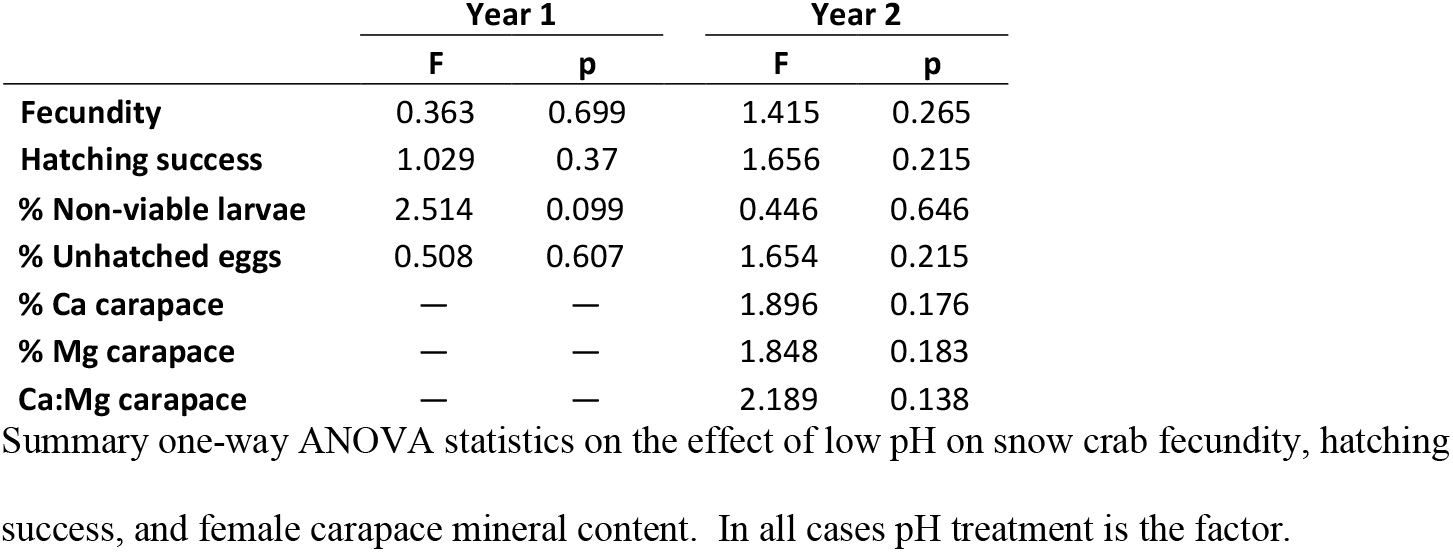
ANOVA result for effects of water pH on snow crab embryos and females.

|  | Year 1 |  | Year 2 |  |
| --- | --- | --- | --- | --- |
|  | F | p | F | p |
| **Fecundity** | 0.363 | 0.699 | 1.415 | 0.265 |
| **Hatching success** | 1.029 | 0.37 | 1.656 | 0.215 |
| % Non-viable larvae | 2.514 | 0.099 | 0.446 | 0.646 |
| % Unhatched eggs | 0.508 | 0.607 | 1.654 | 0.215 |
| % Ca carapace | — | — | 1.896 | 0.176 |
| % Mg carapace | — | — | 1.848 | 0.183 |
| Ca:Mg carapace | — | — | 2.189 | 0.138 |

Summary one-way ANOVA statistics on the effect of low pH on snow crab fecundity, hatching success, and female carapace mineral content. In all cases pH treatment is the factor.

## Discussion

In this study, snow crabs were held for two years over two brooding cycles at low pH with no detectible effects on embryo development, larval hatching, or female calcification. This demonstrates that both the direct effects of low pH and the maternal effects (i.e. carryover effects from embryogenesis) on embryos are negligible. Further, this suggests that the mature females are able to easily cope with the physiological demands of low pH, at least when fed *ad libitum.* We concluded that snow crab may be highly resilient in the face of changing oceanic carbonate chemistry, although other life-history stages not examined may be vulnerable.

There were no detectible direct effects of low pH on embryos in this experiment; rate of development, embryo morphometrics, and survival/hatching success were similar across all treatments. Direct effects of OA on embryos are relatively rare in decapods; a fair number of species show no direct negative effects on measured parameters including the fiddler crab *Leptuca thayeri* [55], the Norway lobster *Nephrops norvegicus* [56,57], and the spider crab *Hyas araneus* [32]. In other species low pH has minor effects on parameters, such as hatch timing or duration, which are unlikely to have a strong effect on overall offspring fitness, without affecting embryo development or survival [12,58]. To our knowledge, in only one species have substantial direct negative effects of OA on embryo development and hatching success been detected: the stone crab, *Menippe mercenaria* [11]. Most decapods carry embryos packed tightly together on pleopods and must regularly aerate the clutch to prevent it from becoming hypoxic; however, in between aerations the dissolved oxygen drops in the egg mass [59] and, presumably, the CO_2_ increases commensurately. In blue king crab, *Paralithodes platypus*, O_2_ level varies with location in the egg mass with eggs in the center experiencing O_2_ saturations as low as 50% [59]. Thus the embryo stages of crabs likely need to be well adapted to substantial variance in pCO_2_ levels and this may explain why this life history stage appears to be highly tolerant of OA conditions across many taxa.

Not only were no direct effects observed, but there were no negative carryover effects of high pCO_2_ from oogenesis (also known as maternal effects) on snow crab embryos. Carryover effects, when exposure to a stressor at one life history stage affects fitness at a subsequent stage, are relatively common in OA work, although because the experiments take longer there are fewer examples. In Tanner crabs, exposure to high pCO_2_ during oogenesis had substantial negative carryover effect on both embryos [12] and the subsequent larvae [47]. High pCO_2_ causes changes to gene expression and disrupts oocyte development in Chinese mitten crabs, *Eriocheir sinensis* suggesting direct effects on oogenesis could be a mechanism for these carryover effect [60]. Alternatively, high pCO_2_ could simply increase the energy requirement to maintain homeostasis and thus decrease the energy available for other processes [61] including reproduction. Regardless, the lack of an effect in snow crabs, both in terms of oogenesis and in terms of shell maintenance (i.e. Ca and Mg content), show that this species has the physiological and energetic plasticity to maintain somatic and reproductive processes under hypercapnic stress. That snow crab larvae receive positive carryover effects from OA exposure during both oogenesis and embryogenesis [53] suggests that physiological plasticity is a major contributor.

The difference in the response to high pCO_2_ between Tanner and snow crabs is striking. These two species are congenators, have overlapping distributions in Bering Sea [62], and are able to produce fertile hybrids [63,64]. However, OA reduces hatching success in Tanner crab by more than 70%, reduces calcium content in larvae, increases mortality of both adults and juveniles, reduces juvenile growth, increases hemocyte mortality, and causes both internal and external dissolution of the adult carapace [12,21,23,46,47,65]. Although fewer studies have been done on snow crab, in the comparable studies that have been done (including this one) no significant negative effects have been observed. This is not unprecedented; different populations of the same crab species, *Hyas araneus*, show a different response to high pCO_2_ [24,66] so different species having different responses is not overly surprising. However, the strong difference in the responses and the similarity of the species opens up the opportunity to explore how their physiological differences contribute to their sensitivity/tolerance of high pCO_2_. Future work should focus on examining how high pCO_2_ changes blood chemistry, respiration rate, and gene expression in both species.

It is, in all honesty, a nice change for these authors to be able to report relatively good news in regards to how high pCO_2_ will affect a commercial crab species. With the data currently available we are cautiously optimistic that snow crab are likely to prove resilient in the face of changing oceanic carbonate chemistry. Future work, however, should still be conducted as different life-history stages may respond very differently to OA; larval American lobsters (*Homarus americanus)* are highly resilient whereas juveniles are very sensitive [67] and European lobster (*Homarus gammarus*) have both sensitive and resilient larval stages and a very sensitive juvenile stage [68,69]. Thus, studies examining all larval stages as well the juveniles stage of snow crab should be performed before it can be concluded that the species as a whole will be resistant to OA. Finally, as pCO_2_ in the oceans increases, temperatures are also projected to increase so experiments examining the separate and interactive effects of pCO_2_, temperature, and hypoxia on snow crab are also called for as these can alter the effects of OA on species [66,70,71].

## Acknowledgments

We thank the RACE Groundfish and Shellfish Assessment Programs of the NOAA Fisheries Alaska Fisheries Science Center and the crews of the F/V Alaska Knight and F/V Arcturus for their assistance in securing crabs used in this study. We thank N. Gabriel, N. Sisson, A. Conrad (née Bateman), and staff of the Kodiak Laboratory for help performing the experiments. Previous versions of this paper were improved by comments from P. McElhany. The findings and conclusions in the paper are those of the authors and do not necessarily represent the views of the National Marine Fisheries Service, NOAA. Reference to trade names or commercial firms does not imply endorsement by the National Marine Fisheries Service, NOAA.

